# *Alk*—*Fam150b* (augmentor α) expression in the paraventricular nucleus of the mouse hypothalamus at molecular resolution, and its sensitivity to acute stress

**DOI:** 10.1101/2025.05.30.656990

**Authors:** Laurent Gueissaz, Spyros Sideromenos, Evgenii O. Tretiakov, Robert Schnell, Tibor Harkany

## Abstract

Augmentor α (*Fam150b*) action on the ALK receptor (*Alk*) has gained significance as a hypothalamic signaling pathway with relevance to the control of food intake and energy homeostasis. In contrast, much less is known about the sensitivity of *Fam150b*-*Alk* expression and signaling upon noxious challenges. In this regard, acute stress is of particular interest because augmentor α, released from afferents of the food intake circuit of the arcuate hypothalamus in the paraventricular hypothalamus (PVN), could link stress-induced changes in food consumption. Nevertheless, conflicting data exist on whether *Fam150b* mRNA is expressed in the PVN. Here, we combined single-cell RNA-seq and multiplexed *in situ* hybridization to demonstrate that both *Fam150b* and *Alk* are expressed in the PVN of adult mice, including corticotropin-releasing hormone (CRH)-containing neurons. As such, a dichotomy of CRH neurons is present through their mutually exclusive expression of either *Fam150b* or *Scgn* (secretagogin). *Fam150b* and *Alk* were not co-expressed. When inducing inflammation-associated stress, *Fam150b* but not *Alk* mRNA expression increased in a mifepristone-sensitive manner, implying regulation by peripheral glucocorticoid feedback. We suggest that augmentor α-ALK signaling could underpin, at least partly, stress-induced changes in feeding and the control of body weight.

## Introduction

According to both experimental and human studies^1^, stress has profound effects on eating behaviors. While acute stress restricts food intake, particularly through changing the sympathetic tone innervating the adrenal medullary system, chronic stressors can increase food intake and lead to obesity through the sustained elevation of circulating cortisol levels^1^. The neurobiological basis of this interaction rests in the hypothalamus, where the territories enriched in stress-responsive and appetite-regulating neurons are positioned proximally in the paraventricular (PVN) and arcuate nuclei (ARC)^2–5^, respectively. It is classically accepted that the stress-induced activation of corticotropin-releasing hormone (*Crh*)^+^ neurons in the PVN, and the downstream activation of the hypothalamus-pituitary-adrenal (HPA) axis^6^, mobilize energy stores to release adipocyte-derived (e.g., leptin, ghrelin) and pancreatic (insulin, glucagon) hormones to feed-back regulate the activity of agouti-related peptide (*Agrp*)^+^ and proopiomelanocortin (*Pomc*)^+^ neurons of the ARC^7–9^. *Agrp*^+^ and *Pomc*^+^ neurons produce antagonistic output with the former stimulating (orexinergic) and the latter blunting (anorexinergic) food intake^102,11,12^. This feedback loop then closes with ARC neurons providing abundant afferents to the PVN either directly^13–15^ or indirectly through GABA relay neurons of the bed nucleus of the stria terminals^16^. Thus, the refined and hierarchical neurocircuit organization of the hypothalamus allows for the dynamic, precisely timed transduction of sensory signals to a metabolic code through peripheral, long-range modulators.

Whereas the neurocircuit layout linking the ARC and PVN is well established, the molecular mediators affecting the interplay of stress-activated *Crh*^+^ and *Agrp*^+^/*Pomc*^+^ neurons are still increasing in number and significance in physiological *vs*. pathobiological states. Firstly, *Agrp*^+^ neurons are GABAergic, and can directly inhibit *Pomc*^+^ neurons through local axon collaterals in the ARC. When releasing AgRP in the PVN, the physiological sign of action is similarly inhibitory because AgRP is an antagonist at melanocortin 3/4 receptors (MC3/4Rs)^17^. Alternatively, *Agrp*^+^ neurons can release neuropeptide Y (*Npy*) in the PVN, which binds Y1^18^ and Y5 receptors^19^ to increase appetite^20,21^ and to modulate sympathetic output to lessen brown adipose tissue thermogenesis^22^. Secondly, *Pomc*^+^ neurons are in a large part glutamatergic (*Vglut2*^+^) with α-melanocyte stimulating hormone (αMSH) being a critical determinant of reduced appetite^23^ when binding to stimulatory Gα_s_-coupled MC3/4Rs^24^ in the PVN. Although the antagonism of αMSH *vs*. NPY/AgRP is considered as the prototypic signaling mechanism to tune PVN neurons, the cellular identity of the postsynaptic cell populations remained ambiguous until single-cell RNA-seq data revealed their receptor repertoires and inferred the synaptic wiring of PVN neurons^25–27^.

More recently, the anaplastic lymphoma kinase (ALK/*Alk*) emerged as an additional receptor whose expression and function in hypothalamic neurocircuits could have implications for energy expenditure. ALK is a receptor tyrosine kinase (RTK), and belongs to the same subfamily as the leukocyte receptor tyrosine kinase (LTK)^28,29^. The first association of ALK and synaptic neurotransmission was obtained when using *Alk* null mice^30^. Soon after, a genome-wide association study linked a variant of ALK to thinness, and showed that ALK loss-of-function reduces triglyceride levels in *Drosophila*^31^. Notably, *Alk*^-/-^ mice had lean body mass due to elevated sympathetic activity and reduced adipose depots. Subsequent conditional and cell-type-specific *Alk* deletion^31^, particularly in the PVN, reinforced its relevance for leanness. However, neither the identity nor potential heterogeneity of neurons that express *Alk* in the PVN, if any, is unequivocally clarified. Likewise, if *Alk* expression is sensitive to acute stress, thus potentially modulating stress-induced body weight changes, remains unexplored.

Augmentor α (AUGα/*Fam150b*) and β (AUGβ/*Fam150a*)^32^ are secreted high-affinity ligands for LTK, with AUGα being particularly efficacious to also induce the phosphorylation of ALK^33,34^. *Augα*^-/-^ and/or *Augβ*^-/-^ mice are resistant to age-related weight gain induced by a high-fat diet^35^. This is because *Augα* knock-out raises norepinephrine levels for a heightened sympathetic tone, reduces the weight of white adipose tissues, and drives thermogenesis, fat oxidation, and energy expenditure^4^. Within the hypothalamus, AUGα^+^ neurons include non-overlapping *Agrp*^+^ and *Crh*^+^ cell cohorts^35^, as suggested by single-cell RNA-seq, with *Fam150b* expression increased upon fasting in *Agrp*^+^ neurons of the ARC^35^. A single-cell RNA-seq study even subdivided *Crh*^+^ neurons into *Scgn*^+^ and *Fam150b*^+^ subclusters^25^. While the subdivision of *Crh*^+^ neurons into stress-responsive *Scgn*^+^ neurons that project to the median eminence (*Crh*^Scgn,stress-on^)^26^ and *Crh^Fam^*^150b^ neurons that might adjust the sympathetic tone carries significant conceptual value, a series of single-cell RNA-seq studies failed to detect *Fam105b* in the PVN^36–38^. This ambiguity curtails how an ALK-AUGα signaling axis could link stress and food intake.

Here, we addressed the identity of neurons that expressed *Alk* and/or *Fam150b* mRNAs in the PVN under physiological conditions, as well as upon acute stress induced by inflammatory pain^39^. When combining single-cell RNA-seq, qPCR in microdissected tissues, and high-resolution multiplexed fluorescence *in situ* hybridization in both female and male mice, we found both *Alk* and *Fam150b* mRNAs in glutamatergic/*Crh*^+^ neurons of the PVN. Stress increased *Fam150b* but not *Alk* expression in a mifepristone-sensitive fashion in *Crh*^+40^/*Fos^+^*^41^ neurons, suggesting increased signaling due to ligand excess and sensitivity to peripheral glucocorticoid feedback. We used MC3/4Rs and opioid receptors (because of their binding of αMSH/adrenocorticotropic hormone (ACTH), and β-endorphin derived from POMC^42^, as well as AgRP itself) as ‘landmarking tools’ to test if *Fam150b* and/or *Alk*-containing neurons in the PVN could serve as post-synaptic partners to either *Pomc*^+^ or *Agrp*^+^/*Npy*^+^ neurons of the ARC. Our results suggest that ALK-AUGα interactions can follow intercellular signaling principles in the PVN, with many *Fam150b*^+^ neurons being likely targets of intrahypothalamic afferents.

## Materials and Methods

### Single-cell RNA-seq analysis

We obtained single-cell RNA sequencing data from three studies on the PVN of adult mice^37,43,44^ (https://harkany-lab.github.io/Gueissaz_2025/03-upset.html#pvn-neurons-from-both-datasets-joined). Initial quality control included filtering the cells based on unique feature counts, unique molecular identifier (UMI) counts, and mitochondrial gene percentage. Data were normalized using *SCTransform*. Variable features were identified using the ‘*vst*’ method selecting 3,000 variable genes.

#### Data integration

For the dataset in Kim *et al*.^37^, we performed reference-based annotation by anchor-based integration with reference data from Romanov *et al.*^44^. Cell-type labels were transferred using *FindTransferAnchors* and *TransferData* functions with 30 dimensions. UMAP embeddings were generated using *RunUMAP* with optimized parameters (n_neighbors = 15-35, min.dist = 0.05-0.8) determined through systematic parameter search by *scDEED*^45^.

#### Subset analysis

PVN neurons were clustered based on hormone and neuropeptide signatures for oxytocin (*Oxt*), vasopressin (*Avp*), somatostatin (*Sst*), *Crh*, and thyrotropin-releasing hormone (*Trh*) across pubertal and adult stages in Seurat v5.1.0 using the ‘*FindNeighbors*’ and ‘*FindClusters*’ functions (**Figure 1a,b**). Co-expression patterns of metabolic (*Alk, Fam150a/b, Mc3r/4r, Lepr, Insr, Lmo4, Irs1/4*), and opioid system-related genes (e.g., *Oprd1, Oprk1, Oprl1, Oprm1, Pcsk1/2, Pdyn, Penk, Pnoc*; **Supporting Figure 1-4**) were assessed for the systematic characterization of gene expression patterns for metabolism-related genes, and their combination with neuropeptide markers in developmentally-distinct neuronal clusters. Intersection analysis for co-expression was performed using the *UpSetR* package (**Figure 1e**). Expression thresholds were determined using the 0.5^th^ percentile of non-zero expression values for each gene.

**Figure 1.**
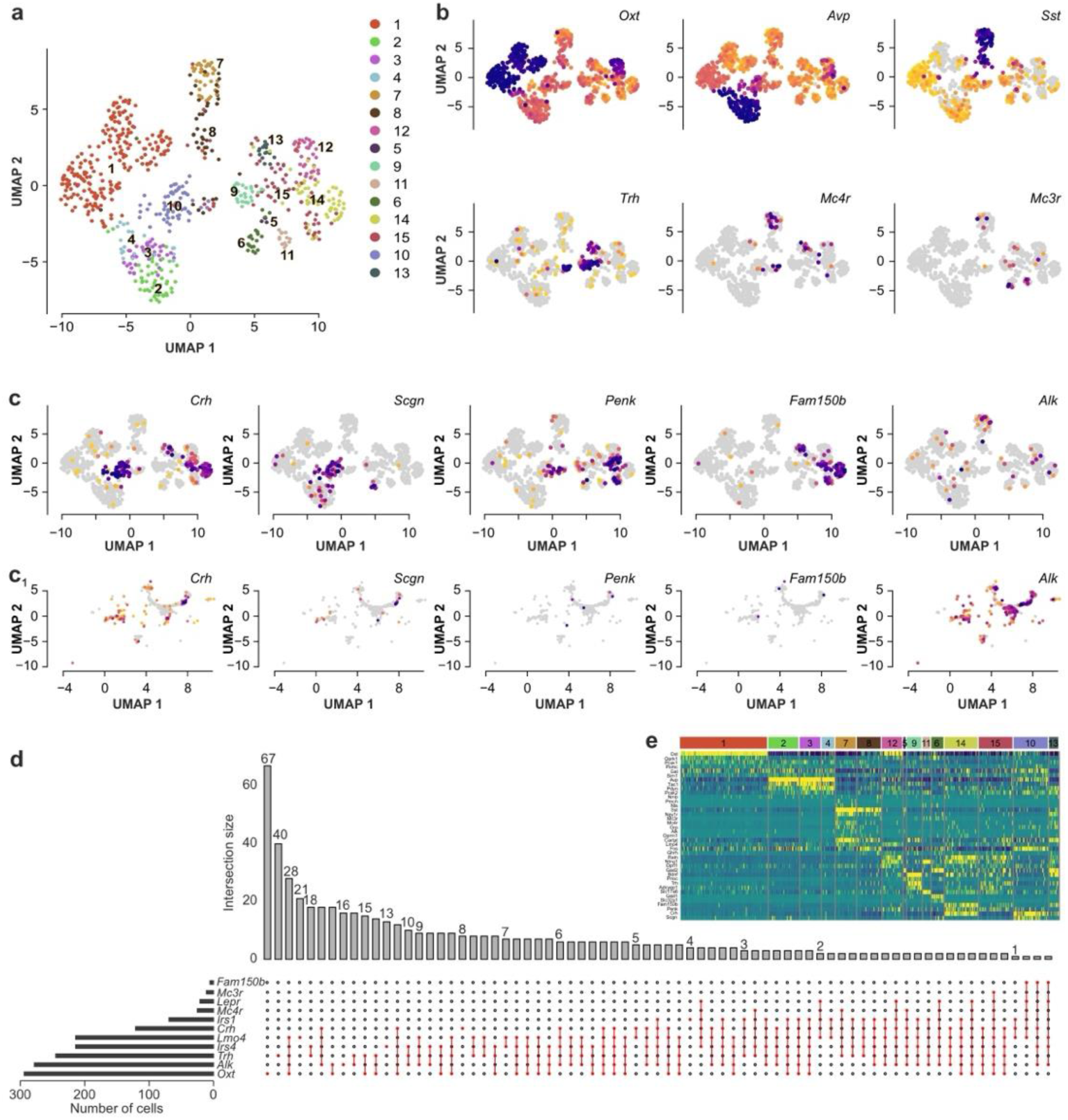
Cellular heterogeneity and gene expression patterns in the PVN. (**a**) Uniform Manifold Approximation and Projection (UMAP) plot for the unsupervised clustering of PVN neurons based on Smart-seq2 single-cell RNA-seq data from Xu *et al*.^25^ used as reference for integration and annotation for 10x-based studies^43,44^. (**b**) UMAP plots for neuropeptides (*Oxt, Avp, Sst, Trh*) and melanocortin receptors (*Mc3r, Mc4r*) across cell clusters. Note the distinct localization of these canonical marker genes to specific cell clusters. (**c**) UMAP plots for the expression of key genes for *Crh* neurons, particularly *Scgn*, *Penk*, *Fam150a*/Augβ, *Fam150b*/Augα, and *Alk* from two reference datasets: (**c**) representing reference dataset used for clustering in (**a**); (**c_1_**) shows expression in postnatal data integrated from single-cell RNA-seq on postnatal day 10 and 23 hypothalami^37,44^. Note the variability in detecting marker genes in *Crh*^+^ neurons with *Fam150a* being almost undetectable, and *Fam150b* only specific for Smart-seq2 data^25^. *Alk* expression was evident in 10x studies^43,44^. (**d**) Co-expression of metabolic genes and cell identity markers in PVN neurons. *Upset plot* visualizing the intersections for select gene combinations was based on integrated 10x single-cell RNA-seq data as above. Horizontal bars to the left show the total number of cells expressing each gene above the 10^th^ percentile threshold. Vertical bars indicate the number of cells expressing specific gene combinations, as per the inter-connected markers below. (**e**) Heat-map for the expression levels of selected neuropeptides, receptors, and signalling molecules across the 15 identified cell clusters in the PVN. Rows represent genes, whereas columns represent clusters. Expression levels were scaled from blue/purple (*low*) to yellow (*high*) to aid the visualization of cluster-specific gene signatures. Expression values in (**b,c,e**) were normalized by total UMI counts scaled to 10,000 molecules and log-transformed with *pseudocount* 1.

#### Data visualization

Visualization was performed using custom *ggplot2*-based functions (v3.5.1), with gene expression features plotted on coordinates in Uniform Manifold Approximation and Projection (UMAP^46^) and *t*-distributed Stochastic Neighbor Embedding (t-SNE)^47^, also allowing the reduction of dimensionality. Statistical summaries of gene expression patterns were generated using the *skimr* package. The analysis pipeline was implemented in *R* (v4.4.1) with key packages including *Seurat* (v5.1.0), *SeuratWrappers*, and *Python 3.8.8* with *scanpy* for data preprocessing. Computational analyses were performed using parallel processing with 8 cores to optimize performance. Code reproducibility was ensured through consistent random seed setting (seed = 42) and the explicit version control of all dependencies.

#### Statistics

Estimation included quantile-based thresholding for gene expression and the systematic evaluation of co-expression patterns. The analysis incorporated quality control metrics including filtering for minimum gene counts, maximum mitochondrial content, and doublet score thresholds^48^.

### Animals and ethical considerations

All experiments were conducted in C57BL/6JRj mice of both sexes at 8-12 weeks of age (Janvier). The sizes of animal groups were specified for each experiment separately. In brief, qPCR was performed on microdissected tissues from *n* = 12 female mice in total, while *n* = 10 female and *n* = 41 male mice were used for molecular neuroanatomy. Animals were randomized, when necessary, yet without blinding. Initially, the animals were group-housed conventionally (12h/12h light/dark cycle, 22-24 °C ambient temperature, and 55% humidity), and allowed ≥7-days of recovery upon arrival to reduce the impact of transport- and environmental change-related stress. Food and water were available *ad libitum*. Experiments on live animals conformed to the 2010/63/EU European Communities Council Directive and were approved by the Austrian Ministry of Science and Research (66.009/0277-WF/V/3b/2017). Effort was directed towards minimizing the number of animals used and their suffering during the experiments. Group sizes conformed to those in the literature^39,44,48^.

### Stress induction and mifepristone treatment

To evaluate the impact of inflammation-associated acute stress on *Alk* and *Fam150b* mRNA expression, mice were randomly assigned to one of three groups (in equal number), that is, control, 30 min (‘30 min’), and 2h survival after stress induction (‘2h’). Stress was induced by 4% paraformaldehyde (PFA) injected subcutaneously in the left hind paw (in a volume of 50 µl)^26^, and confirmed in each case by increased grooming/licking, and reduced mobility. Control mice received physiological saline (50 µl) instead.

To assess if changes in *Fam150b* expression in the PVN were induced by glucocorticoid receptor (GR) activation, mifepristone, a partial GR agonist^49,50^, was used prior to stress induction. Briefly, male mice were randomly assigned to control (*n* = 6), stress ‘30 min’ (*n* = 7), or mifepristone + stress ‘30 min’ (*n* = 7) groups. Control and stress-only animals received an intraperitoneal injection of vehicle (80% saline, 10% dimethyl-sulfoxide, 10% Cremophor) only, at a volume of 10 µl/g bodyweight. Mifepristone was administered at 50 mg/kg bodyweight^51^. To allow for drug action, mice rested for 60 min before receiving a subcutaneous injection of 4% PFA or vehicle as above. Animals were processed 30 min after stress induction.

### Tissue collection

At the specified time-points (‘30 min’ *vs*. ‘2h’), animals were deeply anesthetized (5% isoflurane in 1L airflow/min), their brains rapidly removed, immersed in optimal cutting medium (OCT; Sakura) in plastic molds, and flash-frozen on a combination of liquid N_2_ and ice. Brains were stored at -80°C until use.

### qPCR

Fresh-frozen brains were coronally sectioned at 400-µm thickness on a cryostat microtome (CryoStar NX70, Thermo Fisher). The PVN at both sides was collected from two consecutive sections (from bregma -0.46 mm to -1.26 mm) using a pre-chilled scalpel. Tissues were lysed, RNA isolated using the Aurum Total RNA kit (Bio-Rad), with concentrations determined on a NanoDrop 2000 (Thermo Fisher), and adjusted to transcribe 150 ng/µl into cDNA by using a high-capacity cDNA reverse transcription kit (Applied Biosystems). cDNA was amplified using SYBR green (BioRad) on a CFX Connect Real Time System (Bio-Rad) with mouse-specific primers (Eurofins) as follows: *Crh* (*forward*: ATC TCT CTG GAT CTC ACC TTC C, *reverse*: CCC GAT AAT CTC CAT CAG TTT CC), *Alk* (*forward*: ACT GAC ATC CTC GCT TCT GAA, *reverse*: ATA CGT TTC CTC TCA AAA CCC C), *Fam150b* (*forward*: AGG TTG CTA GTT GAG CTG GTC, *reverse*: CTC CTC TTG GTC TGC CCC ATA) and *Fos* (*forward*: TGG TGA AGA CCG TGT CAG GA, *reverse*: CCT TCG GAT TCT CCG TTT CTC T). TATA-binding protein (*Tbp*) served as internal control (*forward*: CCT TGT ACC CTT CAC CAA TGAC, *reverse*: ACA GCC AAG ATT CAC GGT AGA).

### *In situ* hybridization

*In situ* hybridization procedures were optimized and benchmarked using *n* = 6 adult male mice prior to batch-processing all experimental tissues. Briefly, fresh-frozen brains were coronally sectioned at 16-µm thickness on a cryostat and stored at -20°C until use. The HCR RNA-FISH protocol for ‘*fresh frozen tissue sections*’ (Molecular Instruments) was followed. Sections were air-dried for 10 min prior to fixation (ice-cold 4% PFA for 25 min), dehydrated in an ascending ethanol gradient (30%, 50%, 70% and 100%, 5 min each), and dried for 3 min. Probes (*Crh, Slc17a6, Gad2, Fam150b, Alk, Fos, Scgn, Mc4r and Npy1r*) were from Molecular Instruments. Probe combinations included *Crh/Fam150b/Fos, Crh/Fam150b/Scgn, Crh/Alk/Fos, Crh/Scgn/Fos, Crh/Alk/Scgn, Alk/Mc4r, Alk/Npy1r*, *Alk*/*Slc17a6*, *Alk*/*Gad2*, and *Crh/Alk/Fam150b*. Probes and hairpins were used at a concentration of 0.5 µl/100 µl and 2 µl/100 µl, respectively. Nuclei were counterstained with Hoechst 33,342 (1:5,000; Sigma).

### Confocal microscopy and quantification

Imaging was performed on a Zeiss LSM800 laser-scanning microscope equipped with a 40x oil objective and line lasers for maximal signal separation. Images were acquired in the ZEISS ZEN software (v.2.3). Clustering of the cell populations was in ImageJ^52^ using a semi-automatic toolbox for object-based co-localization^53^. Only images in which the PVN could be unambiguously identified were included in the analysis, averaging at *n* =3 images/ brain (both hemispheres) for each marker combination. The PVN was manually encircled, and the cells semi-automatically grouped according to their mRNA expression of interest.

### Statistics

Histochemical data were normalized to the total number of neurons per PVN in each section. Data were expressed as means ± s.d., and analyzed using GraphPad Prism (v.10.2.3 for Windows). Data were statistically evaluated using one-way ANOVA with Fisher’s least significant difference (LSD) test, where appropriate. A *p* value of <0.05 was considered significant. Data in each figure were plotted as bar graphs overlain with individual data points.

## Results

### Molecular identity of *Alk^+^* and/or *Fam150b^+^* neurons in the PVN

Single-cell RNA-seq data were analysed for neurotransmitters (**Figure 1a,b**) and the co-expression of *Alk*, *Fam150b*, *Scgn* (secretagogin; **Figure 1c; Supporting Figure 1**), *Mc3r/Mc4r, Lepr,* and *Isr3/4* (**Figure 1d**). The expression profile of *Lmo4*, a transcriptional regulator expressed in both *Crh*^+^ and thyrotropin-releasing hormone (*Trh*) ^+^ neurons, was also determined and used for metabolic predictions in juvenile/adult wild-type mice (both C57Bl6/N/J and CD1 strains; **Supporting Figure 1**). *Oxt*^+^, *Avp*^+^, *Sst*^+^, *Crh*^+^ (∼16% of PVN cells), and *Trh*^+^ neurons were subclustered (**Figure 1a,b; Supporting Figure 2-4**). *Oxt*^+^ and *Avp*^+^ neurons did not express appreciable levels of marker genes for fast synaptic neurotransmission (**Figure 1e**). *Sst*^+^ neurons were exclusively GABAergic (contained both *Gad1* and *Slc32a1*). In contrast, *Crh*^+^ and *Trh*^+^ neurons were glutamatergic (*Slc17a6^+^*) (**Figure 1e**).

*Alk* was co-expressed in subsets of *Oxt*^+^ > *Trh*^+^ > *Crh*^+^ neurons under physiological conditions (**Figure 1c,c_1_**) in ∼37% of all PVN neurons. *Alk* and *Fam150b* were particularly enriched in those *Crh*^+^ neurons that co-expressed *Scgn* along with *Oprl1* (nociceptin receptor 1; (∼35% of all cells) and *Oprm1* (μ-opioid receptor; ∼19% of all cells; **Figure 1c,c_1_**). Another *Fam150b*^+^ neuronal cluster had lower levels of *Crh* together with *Penk* mRNA transcripts. It is noteworthy that *Fam150b* expression varied in the reference datasets: Xu *et al*.^25^ who used *Agrp*-IRES-Cre mice for sequencing (*see also* Ref.^35^) detected shallow gene expression (**Figure 1c**). In contrast, Kim *et al*.^37^ found both populations of neurosecretory *Crh^+^* neurons to barely contain, if any, *Fam150b* transcripts (**Figure 1c_1_**). The integration of multiple datasets validated the presence of *Alk* and *Fam150b* in the PVN, suggesting a conserved molecular architecture for metabolic signalling. *Alk* and *Fam150b* were not co-expressed (**Figure 1d**). *Mc3/4r* expression marked *Oxt*^+^, and *Trh*^+^ neurons, but not *Crh*^+^ neurons (**Figure 1b,c**), and co-existed with neither *Fam150b* nor *Alk*.

Cumulatively, these data suggest that *Alk* and *Fam150b* expression are cellular features for some PVN neurons, including *Crh*^+^ cells, and allow for intercellular, rather than cell-autonomous, signalling within the PVN and/or in projection areas. Given the lack of reliable commercial antibodies available to date, we localized mRNAs to molecularly characterize either *Alk*^+^ or *Fam150b*^+^ neurons in the PVN itself.

### Methodological considerations for stress-induced neuronal activation

Quantitative real-time PCR was first used to reproduce earlier studies^39^ on stress-induced neuronal sensitization in the PVN. We have empirically selected 30 min and 2h time-points based on our own and others’ findings^39,54,55^. At 30 min, *Fos* mRNA levels increased by ∼5.4-fold upon stress induction (*p* < 0.001; **Figure 2a**). Similarly, *Crh* mRNA expression was elevated relative to controls (*p* = 0.026; **Figure 2b**). At 2h, *Fos* expression approximated the control value (<2-fold; *p* < 0.001; **Figure 2a**) alike the *Crh* mRNA content (*p* > 0.2; **Figure 2b**).

**Figure 2.**
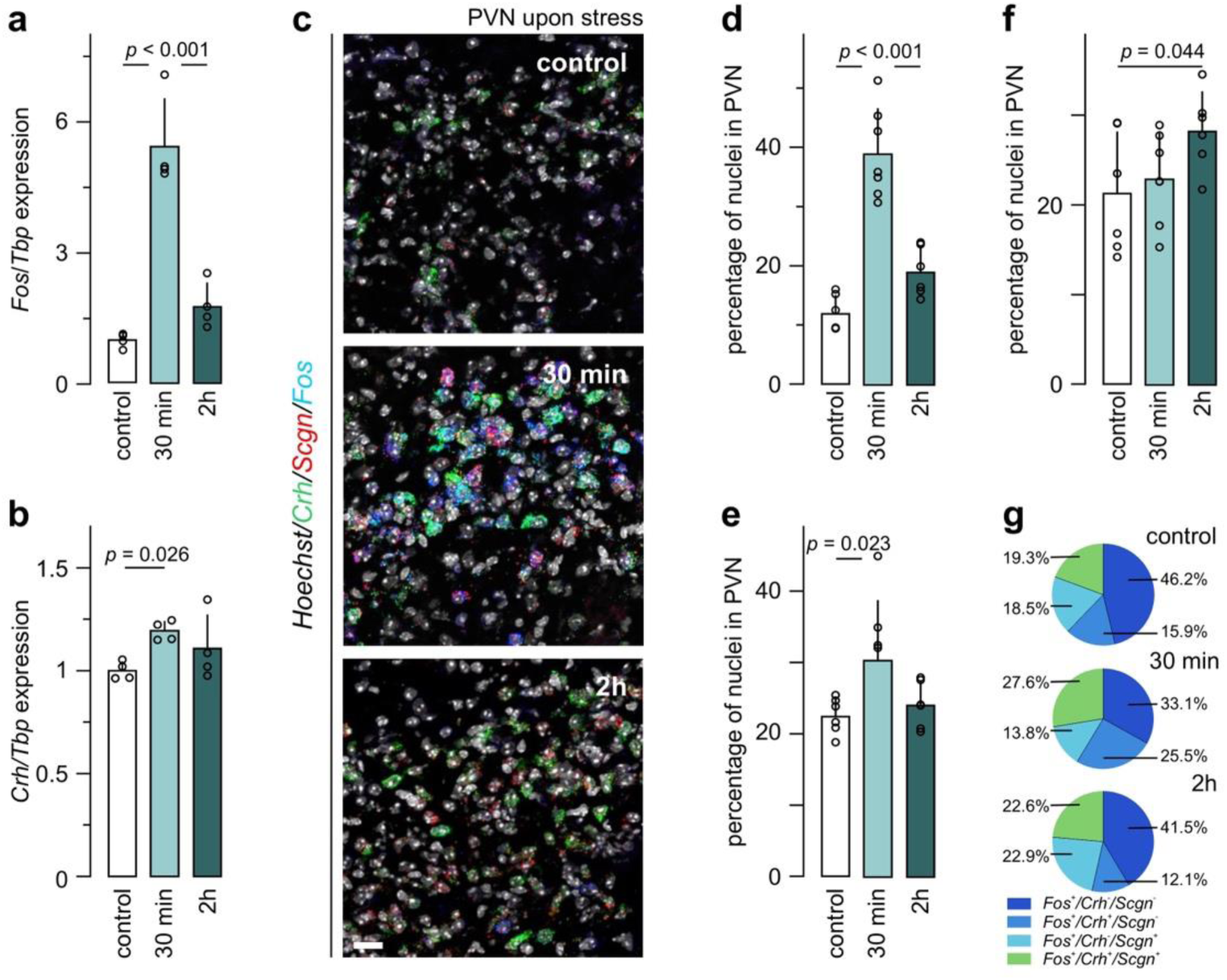
The effect of inflammatory pain on *Fos*, *Crh,* and *Scgn* mRNA expression in the PVN. Quantitative PCR (qPCR) results comparing the impact of acute PFA-induced stress on *Fos* (**a**), and *Crh* (**b**) mRNA expression in the PVN. Equivalent group sizes (*n* = 4) of female adult C57Bl6/N mice were used throughout. Data were normalized to TATA-binding protein (*Tbp*) that had served as a housekeeping standard. Single-molecule fluorescence *in situ* hybridization (**c**) and quantification of *Fos*^+^ (**d**), *Crh*^+^ (**e**), and *Scgn^+^* (**f**) neurons in the PVN under physiological conditions (‘control’) and in animals terminated after acute stress (30 min *vs*. 2h). Data were normalized to the number of nuclei within manually-delineated PVN per section. (**g**) Colocalization coefficient from triple-label experiments within the *Fos^+^* neuronal population. Statistical differences were considered at *p* < 0.05. *Scale bars* = 25 µm.

Next, we validated the qPCR data by multiplexed *in situ* hybridization (**Figure 2c**) and semi-automatic cell counting. The proportion of *Fos*^+^ cells (expressing >2 fluorescent puncta/cell) increased after 30 min from 12.0 ± 3.0% to 39.0 ± 7.6% of all cells in the PVN (*p* < 0.001 *vs*. [control]), with a reduction to 19.0 ± 4.1% after 2h (*p* < 0.001 *vs*. [30 min]; **Figure 2d**). The proportion of *Crh*^+^ neurons increased significantly after 30 min (22.5 ± 2.5% [control] *vs*. 30.4 ± 8.3% [30 min]; *p* = 0.023; **Figure 2e**) but returned to baseline-like levels after 2h (24.1 ± 3.2%). The proportion of *Scgn*^+^ cells, a stable marker of *Crh*^Scgn,stress-on^ neurons^39^, did not change at 30 min, and was only moderately increased 2h after stress induction (21.4 ± 6.8% [control] *vs*. 28.3 ± 5.1% [2h]; *p* = 0.044; **Figure 2f**). This finding is compatible with a slow response in the regulation of secretagogin expression as seen earlier in, e.g., β-cells of the endocrine pancreas^56^.

Next, we asked if the proportion of neurons co-expressing *Crh* and *Fos* had been affected by inflammatory pain-associated stress. At 30 min, we saw a significant increase in dual-labelled neurons (15.9 ± 8.8% [control] *vs*. 25.5 ± 8.5 [30 min]; *p* = 0.048), which returned to baseline by 2h (12.1 ± 6.4; **Figure 2g**). The number of neurons co-labelled for *Crh, Fos,* and *Scgn* also increased after 30 min (19.3 ± 7.4% [control] *vs*. 27.6 ± 4.2% [30 min]; *p* = 0.047), yet tailed off by 2h (22.6 ± 8.6%).

Cumulatively, both molecular and histochemical data using *Fos* expression as a surrogate of neuronal activity confirmed the fast and transient responsiveness of *Crh*^+^ neurons that otherwise stably express *Scgn* to PFA-induced inflammatory pain-associated acute stress.

### *Alk* mRNA expression in the PVN

Under physiological conditions, 33.0 ± 3.7% of PVN cells expressed *Alk* (**Figure 3a,b**). *Alk* expression dominated in glutamatergic (*Slc17a6*^+^) neurons (80.8 ± 6.3% of *Alk*^+^ population; **Figure 3a,c**). Other *Alk*^+^ neurons were GABAergic as judged by the co-existence of *Gad2* (19.2 ± 6.3% of *Alk*^+^ population; **Figure 3a,c**).

**Figure 3.**
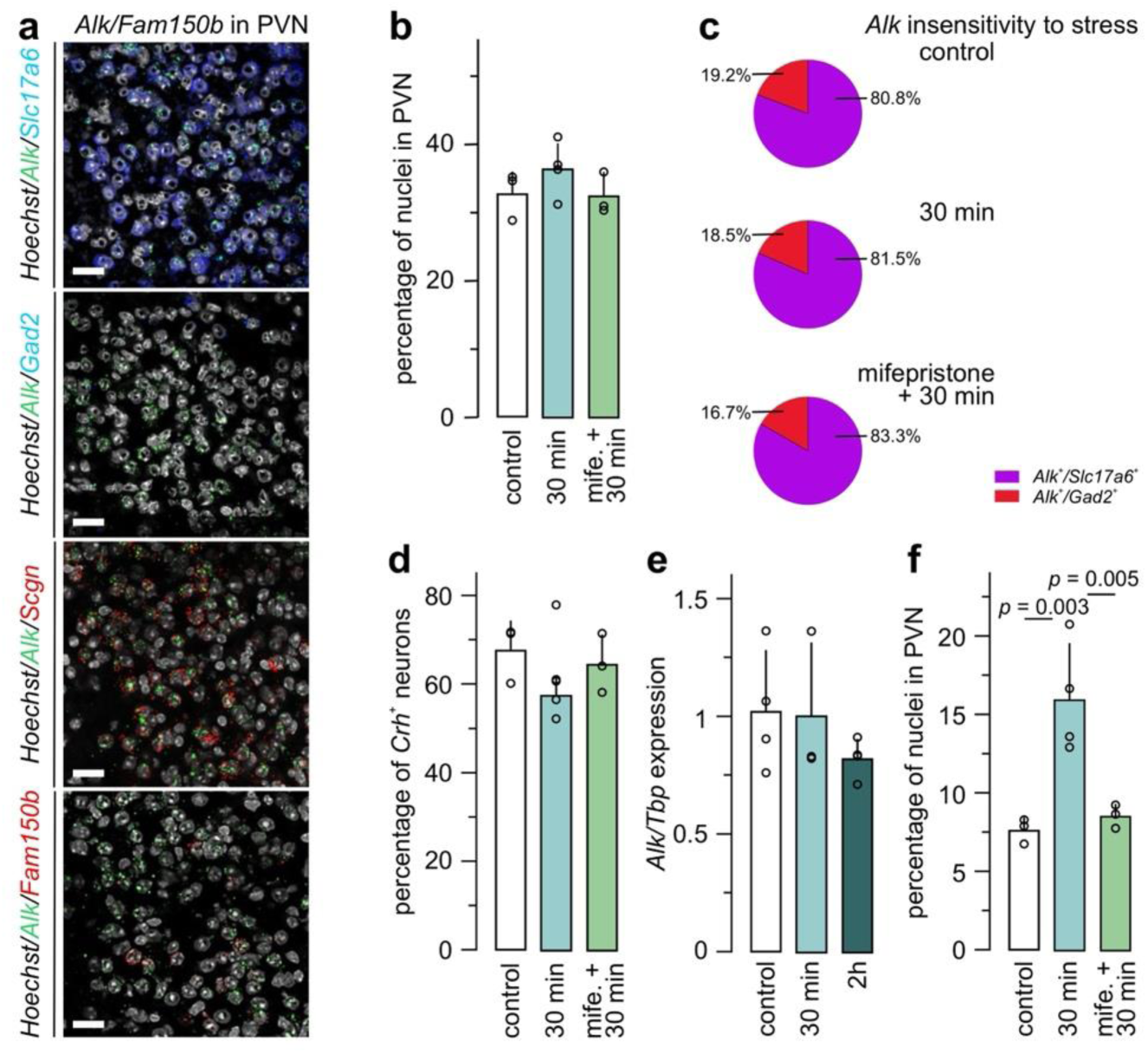
*Alk* mRNA expression in the PVN and its insensitivity to acute inflammation-associated stress. (**a**) Fluorescence multiple-label *in situ* hybridization of the PVN (coronal sections) co-labelled for *Alk with* respectively, *Slc17a6*, *Gad2*, *Scgn* or *Fam150b*. *Scgn* was used a surrogate molecular marker for *Crh*^+^ neurons^39^. No co-localization of *Alk* and *Fam150b* was observed in the PVN (*bottom*). Experiments were conducted in control (vehicle, *n* = 6; all male), PFA-treated (30 min, *n* = 7), and mifepristone + PFA-treated (10 µl/g body weight; *n* = 7) mice. (**b**) Neither stress (30 min) nor mifepristone affected the proportion of *Alk^+^* cells in the PVN. (**c**) The majority of *Alk* mRNA expression was seen in glutamatergic (*Slc17a6*^+^) neurons, with lesser contribution of GABAergic neurons. *Crh*^+^ neurons were sub-grouped and tested for *Alk* mRNA expression. (**d**) The total population size of *Crh^+^/Alk^+^* PVN neurons did not change (control (*n* = 3) *vs*. stress (*n* = 4) *vs.* mifepristone + stress (*n* = 3). (**e**) Quantitative PCR confirmed the lack of change in *Alk* mRNA expression in microdissected PVN samples. Data were normalized to TATA-binding protein (*Tbp*), used as housekeeping standard. (**f**) The proportion of *Fos^+^/Crh^+^/Alk^+^* PVN neurons increased in the stressed (30 min) group. This effect was offset by the mifepristone pre-treatment. Data were normalized to the number of nuclei within manually-delineated PVN in each section. Statistical differences were considered at *p* < 0.05. *Scale bars* = 25 µm.

Histochemical analysis of *Crh*/*Alk* co-expression found that 67.8 ± 6.6% of *Crh*^+^ PVN neurons co-expressed *Alk* mRNA (**Figure 3d**). These data were qualitatively supported by showing *Alk* mRNA in some *Scgn^+^*, but not *Fam150b*^+^, neurons (**Figure 3a**). In addition, *Alk* co-localized with neither *Npy1r* nor *Mc4r* (**Supporting Figure 5a-b_1_**). Overall, we did not only corroborate the single-cell RNA-seq-based molecular subclassification of *Crh*^+^ neurons but also suggest the prevalence of intercellular AUGα-ALK signaling^25^. The segregation of *Alk* and *Mc4r* expression suggests that POMC-derived αMSH and AUGα released from ARC efferents could target spatially non-overlapping *Crh*^+^ cell cohorts in the PVN^25,35^.

### *Alk* mRNA expression upon acute inflammatory pain

Inflammation-associated stress did not alter the population size of *Alk*^+^ neurons in the PVN (36.3 ± 3.9%; **Figure 3b**). Additionally, it did not influence the molecular identity of *Alk*^+^ neurons (for *Slc17a6*: 81.5 ± 8.1% and for *Gad2*: 18.5 ± 8.1% of *Alk*^+^ population; **Figure 3c**); nor affected the proportion of *Crh*^+^ neurons that harbored *Alk* mRNA (57.6 ± 4.1%; **Figure 3d**). These data were confirmed by qPCR from micro-dissected PVN samples, which revealed unchanged *Alk* expression at both 30 min and 2h after PFA injection (**Figure 3e**). In contrast, the proportion of neurons co-labelled for *Fos*/*Crh*/*Alk* significantly increased 30 min after PFA injection (7.6 ± 0.1% [control] *vs.* 16.0 ± 3.5% [30 min]; *p* = 0.003; **Figure 3f**). These data suggest the insensitivity of *Alk* mRNA expression to pain-associated stress.

Next, we used mifepristone to block the cellular effects of systemic glucocorticoid feedback^49,57,58^. Treatment of mice with mifepristone did not alter the size of the PVN *Alk*^+^ population (32.5 ± 2.9%; *p* = 0.27; **Figure 3b**). *Alk* mRNA expression was not affected in *Slc17a6*^+^ (83.3 ± 5.4%; *p* = 0.48; **Figure 3c**), *Gad2*^+^ (16.7 ± 5.4%; *p* = 0.48; **Figure 3c**), and *Crh*^+^ neurons (64.5 ± 6.7; *p* = 0.15; **Figure 3d**) relative to control subjects. The action of mifepristone was quality-controlled through the reduced density of PVN cells triple-labelled for *Fos*/*Crh*/*Alk* (8.5 ± 0.1%), as compared to PFA-exposed subjects (16.0 ± 3.5%; *p* = 0.005; **Figure 3e**). In sum, these data suggest that *Alk* is chiefly present in *Slc17a6*^+^/*Crh*^+^ neurons of the PVN^31^, and its expression is insensitive to inflammation-induced stress acutely.

### *Fam150b* expression in the PVN

Under physiological conditions, 9.6 ± 2.0% of all cells in the PVN contained *Fam150b* mRNA (**Figure 4a,b**). Among the *Crh*^+^ neurons, ∼27% harbored *Fam150b* hybridization signal (**Figure 4c**). *Fam150b* co-localized with neither *Alk* (**Figure 3a**) nor *Scgn* (**Supporting Figure 5c,c_1_**). These data demonstrate that *Fam150b* is expressed in a subset of glutamatergic/*Crh*^+^ neurons in the PVN, that segregate from their *Scgn*^+^ counterparts^25^.

**Figure 4.**
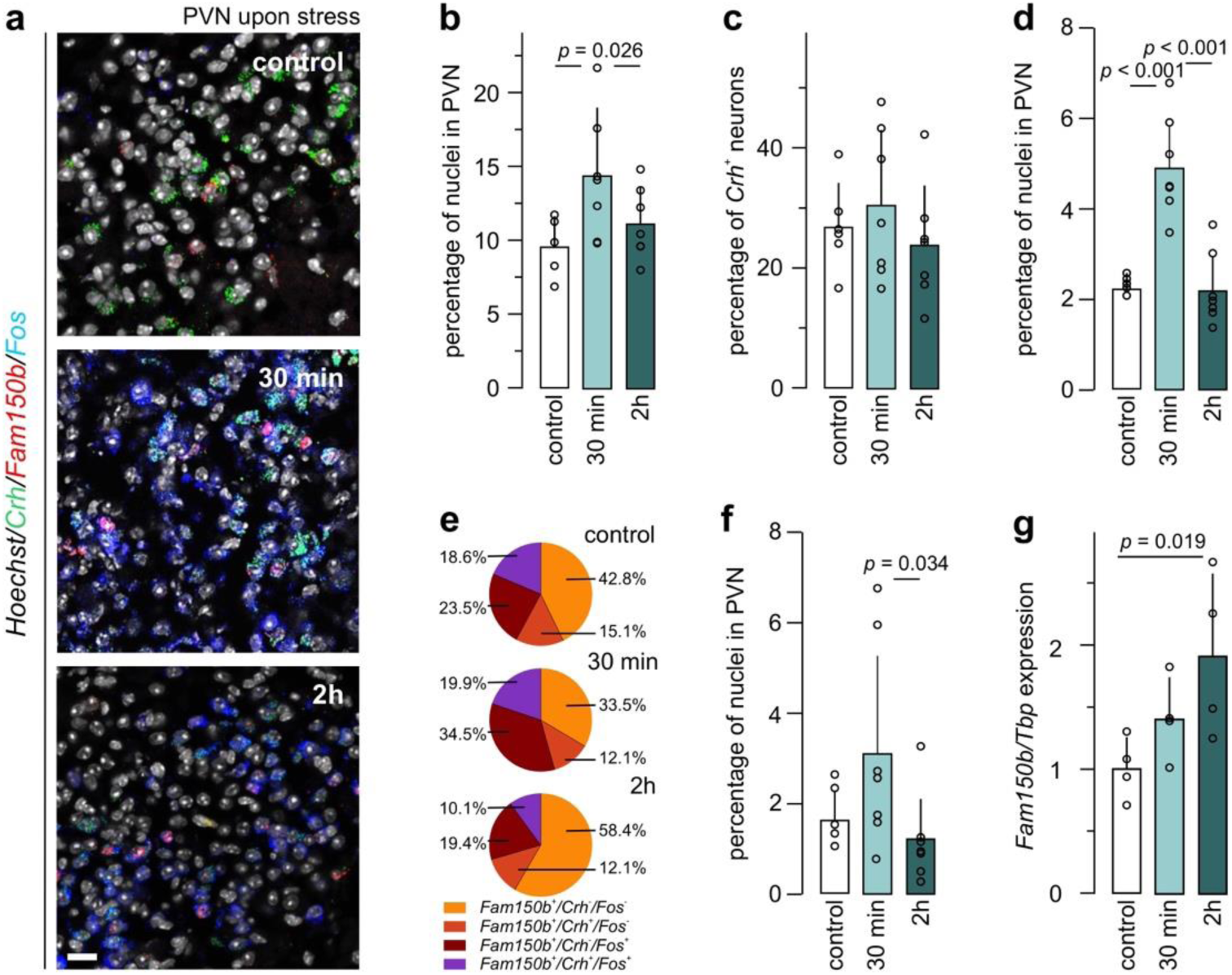
*Fam150b* expression in the PVN upon inflammation-induced acute stress. (**a**) Multiple-label *in situ* hybridization for *Fam150b*, *Crh*, and *Fos* mRNAs in the PVN under control conditions (*n* = 6), and 30 min (*n* = 7) or 2h (*n* = 7) after PFA injection (all adult males). Note the induced expression of *Fos* mRNA in response to stress induction. (**b**) Next, the total number of *Fam150b^+^* neurons in the PVN was determined, revealing a transient increase at 30 min after stress induction. (**c**) Amongst *Crh^+^* PVN neurons, *Fam150b* mRNA expression did not change upon stress induction. (**d**) The proportion of the *Fos*^+^/*Crh*^-^*/Fam150b*^+^population increased in response to stress induction. (**e**) Co-localization coefficient from triple-label experiments within the *Fam150b^+^* neuronal population. (**f**) The proportion of *Fos*^+^/*Crh*^+^*/Fam150b*^+^ neurons increased 30 min and decreased 2h after stress induction, albeit non-significantly relative to controls. Data were normalized to the number of nuclei within the manually delineated PVN for each section. (**g**) Quantitative PCR from microdissected mouse PVN showed a gradual increase in total *Fam150b* mRNA. Data were normalized to TATA-binding protein (*Tbp*), used as a housekeeping standard. Statistical differences were considered at *p* < 0.05. *Scale bars* = 30 µm.

### *Fam150b* expression upon acute inflammatory pain

The population size of *Fam150b^+^* neurons increased to 14.4 ± 4.6% 30 min after PFA injection (*p* = 0.026; **Figure 4a,b**). Additionally, the likelihood of finding *Fos^+^/Fam150b^+^* neurons increased at 30 min (2.2 ± 0.2% [control] *vs.* 4.8 ± 1.1% [30 min]; **Figure 4d**) representing, respectively, 23.5 ± 4.3% and 34.5 ± 5.3% (*p* = 0.003; **Figure 4e**) of the *Fam150b*^+^ population. This change returned to baseline after 2h to 2.1 ± 0.8% (*vs*. [30 min]; *p* < 0.001; **Figure 4d**) of total PVN cells, representing 19.4 ± 7.0% (*vs*. [30 min]; *p* < 0.001; **Figure 4e**) of *Fam150b*^+^ cells. Additionally, the proportion of *Crh*/*Fos*/*Fam150b* triple-labelled neurons in the PVN remained unchanged between control and 30 min, yet slightly decreased 2h after PFA exposure (1.8 ± 0.7% [control] *vs.* 3.1 ± 2.3% [30 min] (*p* = 0.15) and 1.2 ± 1.0% [2h] (*p* = 0.034); **Figure 4f**). These results respectively represented 18.6 ± 3.5%, 19.9 ± 9.2% (*p* > 0.7) and 10.1 ± 6.2% (*p* = 0.034; **Figure 4e**) within the *Fam150b*^+^ population. Next, qPCR was performed on microdissected tissues to corroborate the dynamics of *Fam105b* expression in the PVN. *Fam150b* mRNA content progressively increased, reaching significance at 2h (*n* = 4/group; *p* = 0.019; **Figure 4g**). These data suggest that *Fam150b* in non-*Crh* neurons transiently increased upon inflammation-associated acute stress.

Lastly, we sought to determine if *Fos*/*Fam150b* co-existence was sensitive to the manipulation of systemic glucocorticoid action (alike CRH^59^) by applying mifepristone prior to PFA treatment. PFA injection coincidently increased *Fos, Crh,* and *Fam150b* expression (**Figure 5a**). In this particular experiment, PFA increased the portion of *Fos*^+^ neurons from 10.2 ± 5.7% [control] to 26.4 ± 6.6% [30 min] (*p* < 0.001; **Figure 5b**); *Crh*^+^ neurons from 9.9 ± 3.1% [control] to 17.3 ± 1.8% [30 min] (*p* < 0.001; **Figure 5c**); *Fam150b*^+^ neurons from 5.7 ± 1.6% [control] to 8.9 ± 1.8% [30 min] (*p* = 0.002; **Figure 5d**). Mifepristone prevented the pain-induced increases in any of the mRNAs sampled (*Fos*: 18.9 ± 4.7%; *p* = 0.026; **Figure 5b**; *Crh*: 9.7 ± 2.2%; *p* < 0.001; **Figure 5c**; *Fam150b*: 6.1 ± 1.1%; *p* = 0.003; **Figure 5d**). Additionally, we observed that PFA increased the portion of *Fos/Fam150b* double-positive neurons in the PVN from 1.2 ± 0.7% to 2.9 ± 1.0% (control *vs.* [30 min]; *p* = 0.002; **Figure 5e**), which was prevented by mifepristone i (1.7 ± 0.4%; *p* = 0.008; **Figure 5e**).

**Figure 5.**
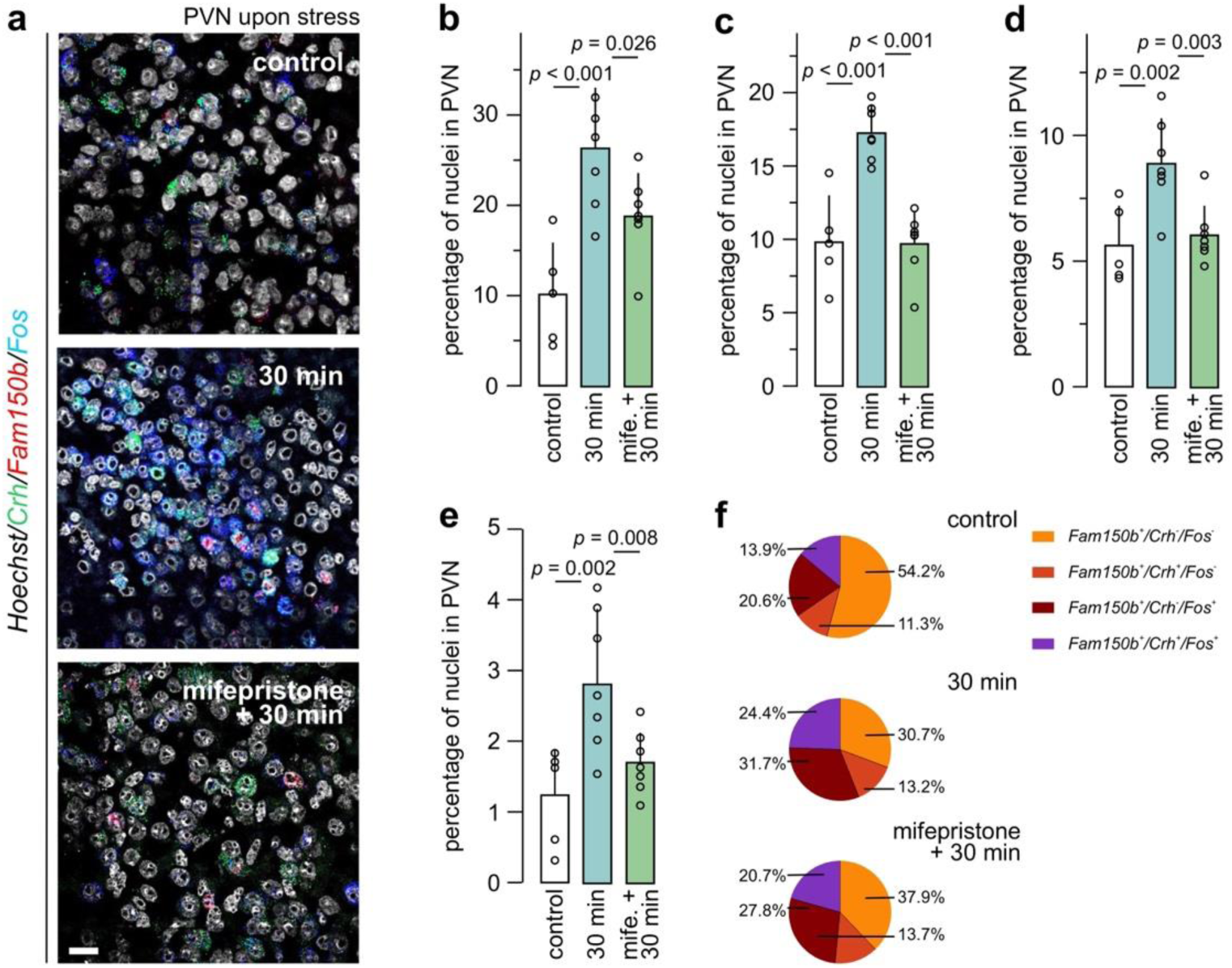
Mifepristone modulates *Fam150b* mRNA expression in the PVN. (**a**) Multiple-label *in situ* hybridization for *Fam150b*, *Crh*, and *Fos* mRNAs in the PVN (adult male mice; *n* = 6 [control], *n* = 7 [acute stress], and *n* = 7 [mifepristone + acute stress]). Acute stress up-regulated *Fos* (**b**), *Crh* (**c**), and *Fam150b* (**d**) mRNAs in a mifepristone-sensitive manner. (**e**) The proportion of the *Fos*^+^/ *Crh*^-^*/ Fam150b*^+^population increased in response to stress induction in a mifepristone-sensitive manner. (**f**) Co-localization coefficient from triple-label experiments within the *Fam150b^+^* neuronal population. Statistical differences were considered at *p* < 0.05. *Scale bars* = 30 µm.

In sum, these data suggest that *Fam150b* expression is sensitive to pain-associated stress in both *Crh*^+^/*Fos*^+^ and non-*Crh/Fos^+^* neurons of the PVN (**Figure 5f**), wherein *Fam150b*^+^*/Crh*^+^ neurons are inferred as a non-overlapping subpopulation with *Alk*^+^*/Crh*^+^ neurons. Thus, our data after mifepristone treatment are consistent with the hypothesis that *Fam150b*^-^/*Scgn*^+^ neurons might be seen as ‘first responders’, while *Fam150b*^+^/*Scgn*^-^ neurons could control sympathetic output instead.

## Discussion

The impact of stress on eating behaviour and weight change is still a topic of scientific discourse. The organizational complexity of the neural networks underpinning the regulation and integration of the origin (modality), severity, and duration of the stress response with the resetting of metabolic set-points remains a major conceptual challenge. As such, experimental designs themselves can be biasing, and even limiting, the experimental outcomes through emphasizing some, while also negating other, variables. Nevertheless, the duration of stress has invariably emerged as a primary factor to affect weight change, with acute and chronic stress reducing and increasing it, respectively^1^. This duality could be, at least in part, due to the participation of either the sympathetic medullary adrenal system to increase catecholamine release (both adrenaline and noradrenaline), with reduced food intake as its consequence. In turn, activation of the HPA axis increases circulating glucocorticoid levels, and provokes weight gain^1^. Besides the brain-periphery axes engaged and provoked, the molecular identity of synaptic mediators within the hypothalamus can also diversify body-wide responses to stress.

Therefore, we have firstly examined the cellular architecture of the ALK-AUGα (*Fam150b*) signaling cassette, which was recently associated with the control of eating behaviour and energy metabolism through modulating PVN activity and output^31,35^. While Fam150b was described as a ligand co-released from ARC efferents to the PVN, neither the cell-type specificity nor stress sensitivity of either ligand (AUGα) or receptor (ALK) expression is known. When merging single-cell RNA-seq and *in situ* hybridization data, we concluded that ∼35% of all cell contained anatomical indices of AUGα-ALK signaling, with the majority of *Alk*^+^ cells being glutamatergic (*Slc17a6*^+^)^35^, rather than GABAergic (*Gad2^+^*; 20% only). This finding is compatible with the predominantly glutamatergic neurochemical signature of PVN neurons.

Many *Crh*^+^ neurons expressed *Alk*, which can be considered as a sub-class marker for this group of neurons. The reason for this is that *Alk* identifies a subset of *Crh*^+^ neurons that (mostly) do not express *Scgn*^25^. Considering that *Scgn* has been causally associated with the initiation of the HPA axis^39^, the subset of *Alk*^+^/*Crh*^+^ neurons could indeed modulate sympathetic output independently as suggested earlier^31^.

*Alk* and *Fam150b* are expressed in non-overlapping populations of neurons in the PVN, thus implying an intercellular ligand-receptor relationship between them. In the ‘classical’ view of the ARC-PVN circuit controlling eating behaviour, AgRP^+^/NPY^+^ neurons in the ARC are upstream to MC4R^+^/NPY1R^+^ neurons in the PVN^17–19^. Genetic studies demonstrate that AUGα produced by AgRP^+^/NPY^+^ neurons is critical to increase weight gain, as much as ALK in the PVN^31,35^. Nevertheless, our neuroanatomy data uncovered a number of peculiarities, including that *Alk* expression decorated PVN neurons distinct from those with either *Mc4r* or *Npy1r* in the PVN. These data are in accord with those of Ahmed *et al*.^35^. Moreover, *Fam150b* itself is expressed in the PVN, including glutamatergic neurons that co-express *Crh* mRNA^25^. Thus, we resolved conflicting single-cell RNA-seq predating our present work^25,37^, and support *bona fide Fam150b* expression in the PVN. We also entertain the possibility that the few local circuits that exist within the PVN could use AUGα to cross-modulate neuronal activity locally. Alternatively, extrahypothalamic efferents originating in the PVN could use AUGα to tune the activity of far-placed target neurons. Taken together, we suggest that the AUGα -ALK ligand/receptor pair could influence weight gain *via* neurocircuits parallel to MC4R^+^ or NPY1R^+^ neurons.

Secondly, we determined how PFA injected into an extremity, and producing inflammatory pain acutely, could impact the expression of either *Alk* or *Fam150b*, or both. We have shown that *Alk* expression in the PVN was insensitive to stress, even if a higher proportion of *Fos*^+^/*Crh*^+^/*Alk*^+^ neurons appeared due to *Fos* activation. In contrast, *Fam150b* mRNA levels increased upon PFA-induced stress, including within the *Fos*^+^/*Fam150b*^+^ neuronal population. These data are congruent with the generally accepted notion on ligand rather than receptor-level regulation of intercellular signaling upon environmental challenges or diseases (sensory modalities, injury models). Notably, *Fam150b* expression was regulated by peripheral glucocorticoids because the inhibition of GRs by mifepristone blunted changes in any constellation of *i*) the total proportion of *Fam150b*^+^ cells, *ii*) *Fos*^+^/*Fam150b*^+^, *iii*) *Fos*^+^/*Crh*^+^/*Fam150b*^+^, and *iv*) *Fos*^+^/*Crh*^+^/*Alk*^+^ cell populations. Thus, *Fam150b* in the PVN is sensitive to stress, in a glucocorticoids-dependent manner.

Our neuroanatomy data support the notion that (at least) two subsets of *Crh*^+^ neurons exist in the PVN, for which conflicting single-cell RNA-seq data have been reported. Even though *Scgn* expression has not been contested, *Fam150b* was often not reported by single-cell transcriptomics. A likely reason for this discrepancy is the low copy number of *Fam150b* mRNAs per cell, regardless of the sequencing method applied (10x Genomics *vs*. Smart-seq; **Figure 1c,c_1_**). Yet, our anatomy data could become conceptually appealing when considering the antagonistic action of CRH and glucocorticoids on eating and metabolism: overexpression of CRH reduces nutrition in fasting mice^60^, whereas corticosterone increases appetite by stimulating AgRP^+^ and inhibiting POMC^+^ neurons in the ARC^61^. A second subset of *Crh*^+^ neurons expressing *Fam150b*, but not *Scgn,* could explain the long-term effect of stress on eating behavior. *Scgn*^+^/*Crh*^+^ neurons could operate as ‘first responders’ to stress, increase CRH release at the median eminence, and ultimately increase circulating corticosteroid levels. In contrast, *Crh*^+^/*Fam150b*^+^ neurons could alter sympathetic outflow upon stress, with their increased activity inferred from the increased presence of *Fos*. Even though we do not yet know the precise site(s) of Fam150b release, we hypothesize that it either provides feed-forward stimulation to *Alk*^+^ neurons locally in the PVN, or at postsynaptic targets of long-range afferents. Another intriguing observation is the presence of *Fam150b* in non-*Crh* neurons, which suggests broader roles for AUGα-ALK signaling than previously thought. Overall, we suggest that the sensitivity of AUGα-ALK signaling in non-overlapping neuronal subsets in the PVN to stress could be significant to diversify the cellular control of metabolic readiness in fight/flight situations.

## Supporting information

Supportive Figures

## Acknowledgements

The authors T. Hökfelt for discussion and feedback, and I. Pitzen for laboratory assistance. This work was supported by the Swedish Research Council (2023-03058; T.H.); Novo Nordisk Foundation (NNF23OC0084476; T.H.); Hjärnfonden (FO2022-0300; T.H.), European Research Council (FOODFORLIFE, 2020-AdG-101021016; T.H.), the Austrian Science Fund (COE-16B, T.H.) and intramural funds of the Medical Neuroscience Cluster of the Medical University of Vienna (2021/1, T.H.).

## Author contributions

T.H. conceived the project; S.S., R.S., and T.H. designed experiments; T.H. procured funding; L.G. and S.S. performed experiments and analyzed data; E.O.T. performed data integration and the analysis of single-cell RNA-seq data; L.G. ant T.H. drafted the manuscript. All authors have proofread and updated earlier versions of this report.

## Data availability

All data relevant to this report appear in the published dataset.

## Code availability

The code developed and used in this report has been published at https://harkany-lab.github.io/Gueissaz_2025/03-upset.html#pvn-neurons-from-both-datasets-joined

## Supporting Figures and their Legends

**Supporting Figure 1.**
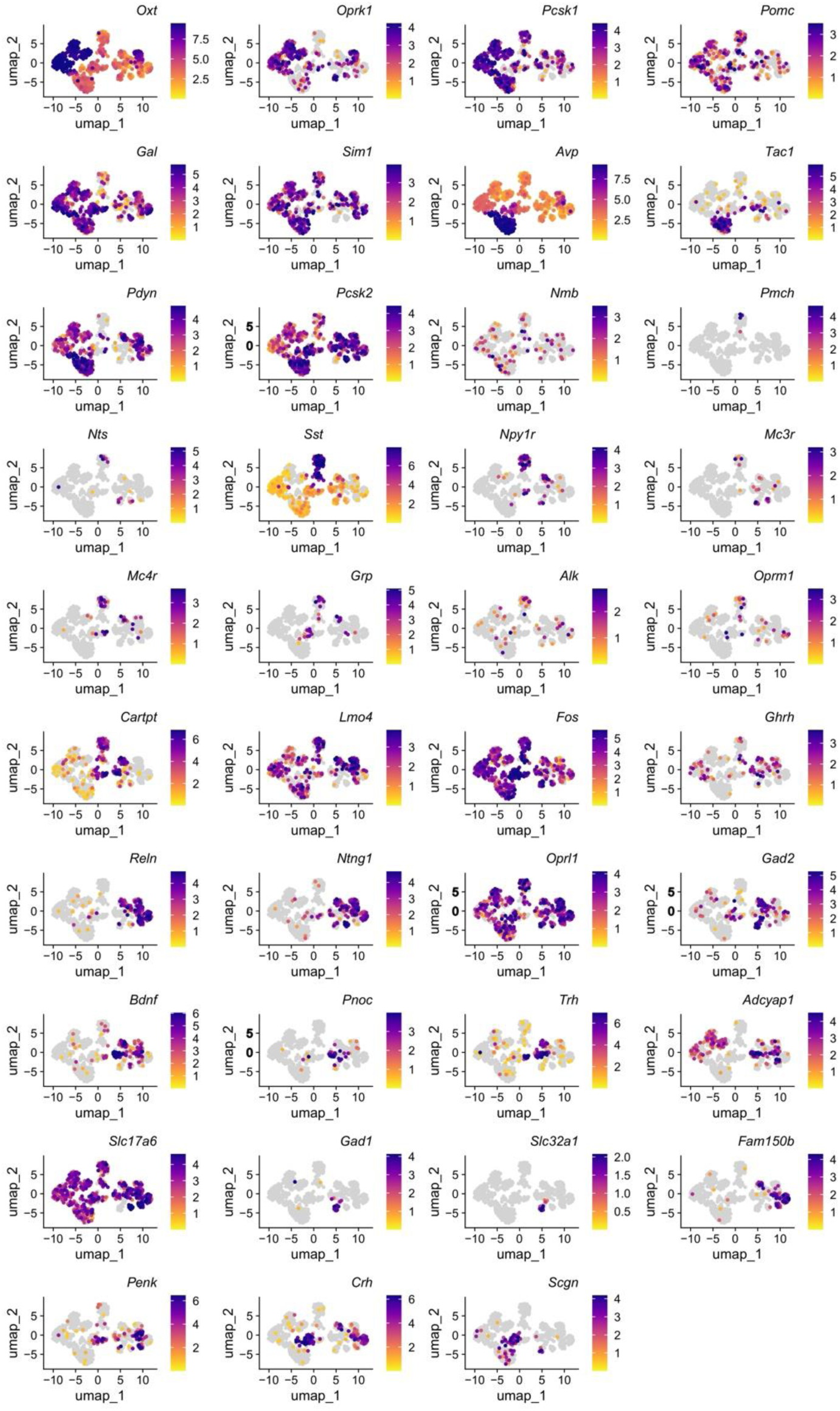
Expression landscape of common neuropeptides and receptors in the mouse PVN. UMAP plots depict normalized expression levels for selected genes in all neurons retrieved from a reference single-cell RNA-seq dataset (Smart-seq2)^25^.

**Supporting Figure 2.**
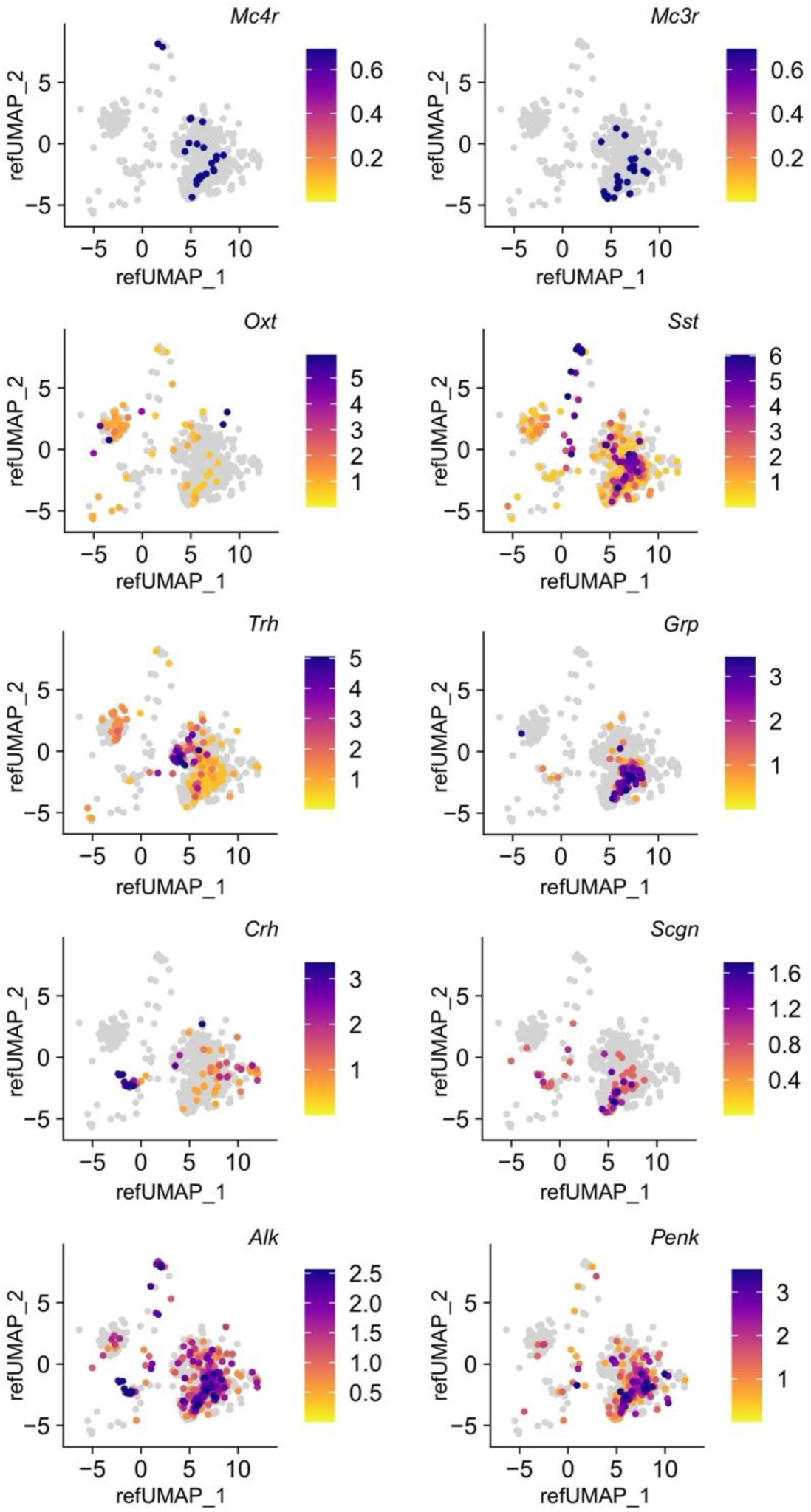
Focused expression of marker and regulatory genes. UMAP plots for the expression of *Mc4r, Mc3r, Oxt, Sst, Trh, Grp, Crh, Scgn, Alk,* and *Penk* from 10x data^43^ mapped onto a reference UMAP from Xu *et al.*^25^.

**Supporting Figure 3.**
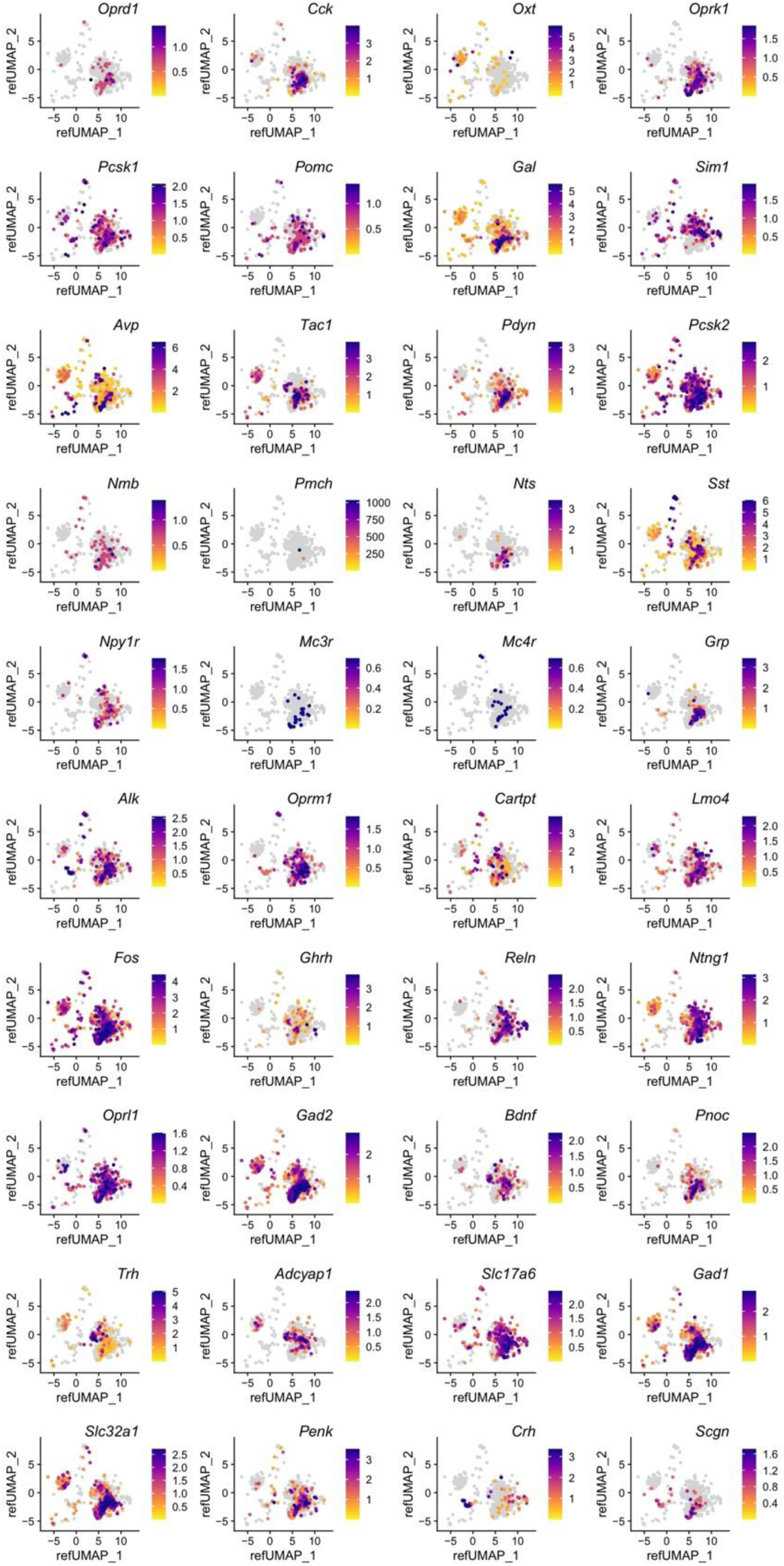
Expression landscape of neuropeptides and with 10x data mapped onto an integrated Smart-seq2 reference. UMAP plots show the expression of genes selected from 10x data^43^ and mapped onto an integrated reference UMAP from Xu *et al.*^25^.

**Supporting Figure 4.**
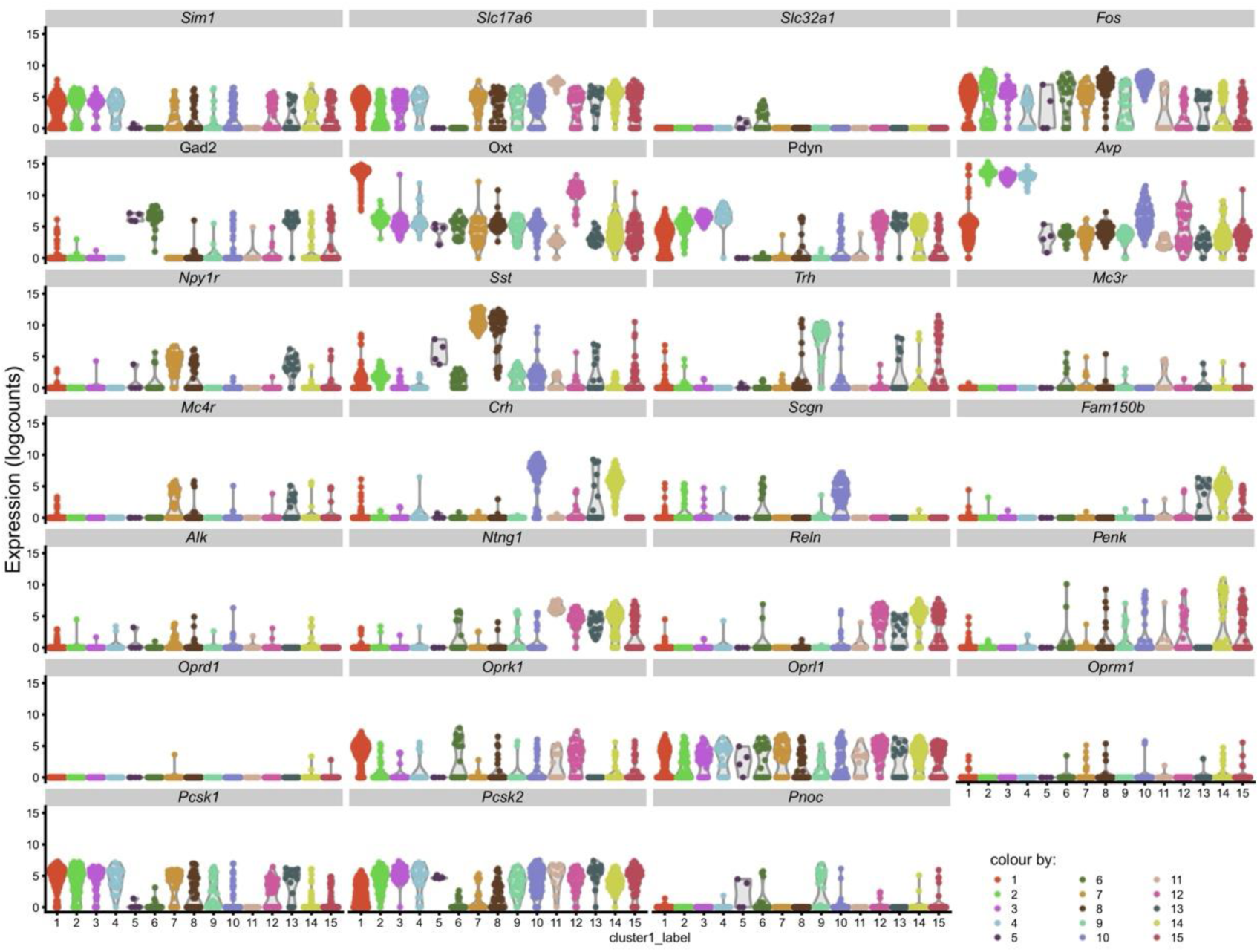
Quantification of gene expression across neuronal clusters in the PVN. Violin plots illustrate the distribution of normalized expression levels (logcounts) for selected genes across cell clusters (not sorted) identified in the PVN by Smart-seq2^25^. Each violin plot shows the density distribution of expression for a specific gene within a given cluster, providing a quantitative comparison of gene expression profiles across neuronal subpopulations.

**Supporting Figure 5.**
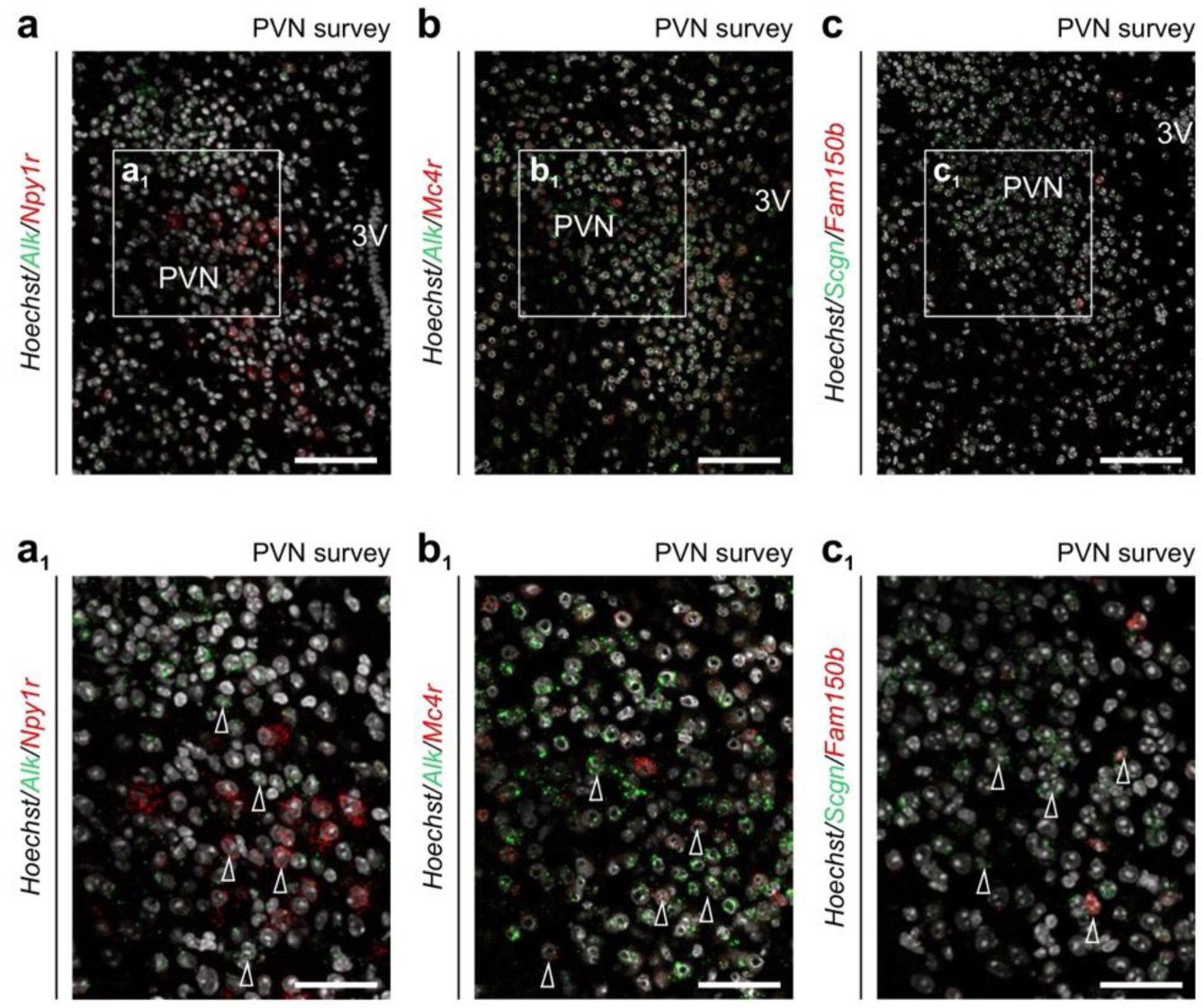
Colocalization of *Alk* and *Fam150b* with other cellular markers in the PVN. Multiple-labelling *in situ* hybridization for *Alk* and either *Npy1r* (**a**) or *Mc4r* (**b**) in the PVN. In both cases, *Alk* did not seem to co-localize with *Npy1r* or *Mc4r*, two receptors implicated in the control of metabolism and feeding behaviors^20–24^. (**c**) Multiple-labelling *in situ* hybridization for *Fam150b* and *Scgn* in the PVN. *Fam150b* and *Scgn* did not co-localize in the PVN, confirming data from single-cell RNA-seq^25^. *Scale bars* = 300 µm (*overviews*) and 30 µm (*insets*).

