## Supplementary material for "*Alk*—*Fam150b* (augmentor α) expression in the paraventricular nucleus of the mouse hypothalamus at molecular resolution, and its sensitivity to acute stress": Supportive Figures

### Supporting Figures and their Legends

Gueissaz *et al*. - Supporting Figure 1

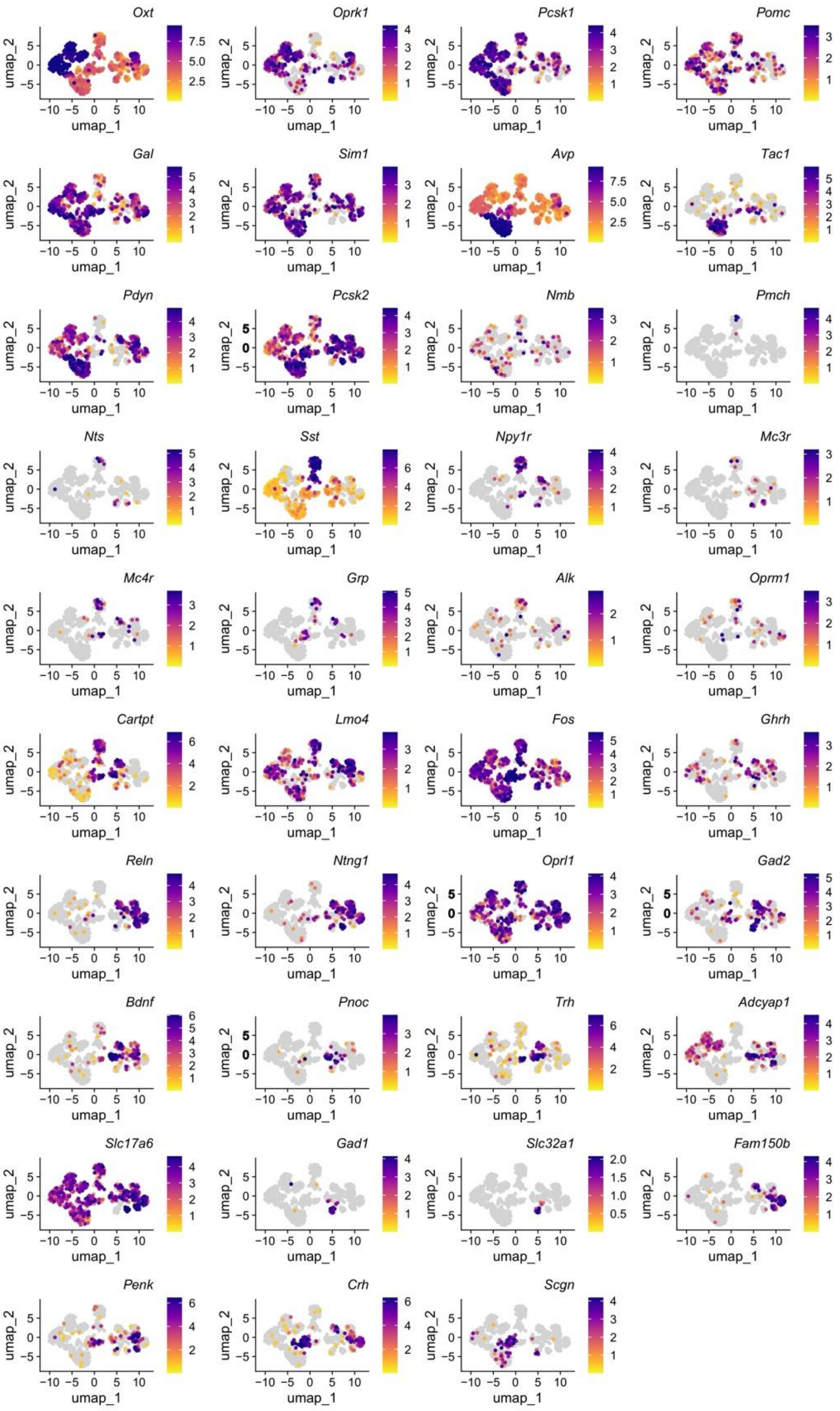


**Supporting Figure 1. Expression landscape of common neuropeptides and receptors in the mouse PVN.** UMAP plots depict normalized expression levels for selected genes in all neurons retrieved from a reference single-cell RNA-seq dataset (Smart-seq2)[25].

Gueissaz *et al.* - Supporting Figure 2

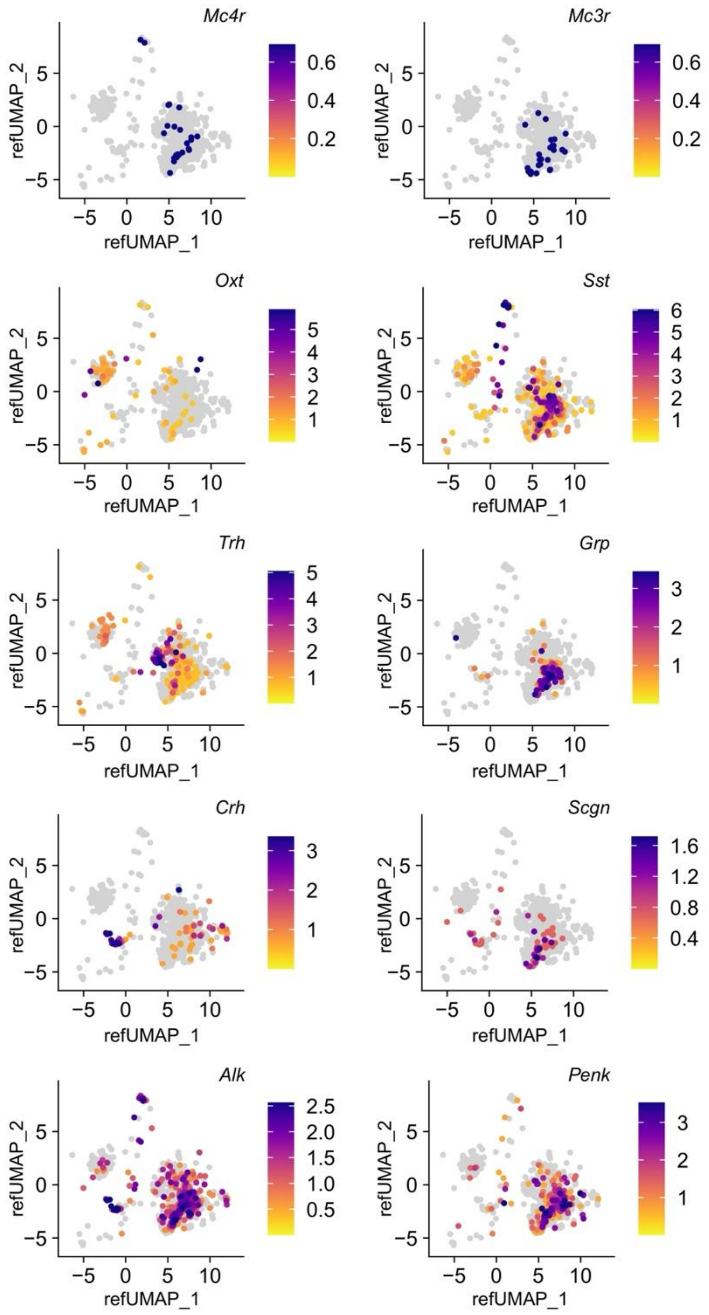


**Supporting Figure 2. Focused expression of marker and regulatory genes.** UMAP plots for the expression of *Mc4r, Mc3r, Oxt, Sst, Trh, Grp, Crh, Scgn, Alk,* and *Penk* from 10x data[43] mapped onto a reference UMAP from Xu *et al.*[25].

Gueissaz *et al*. - Supporting Figure 3

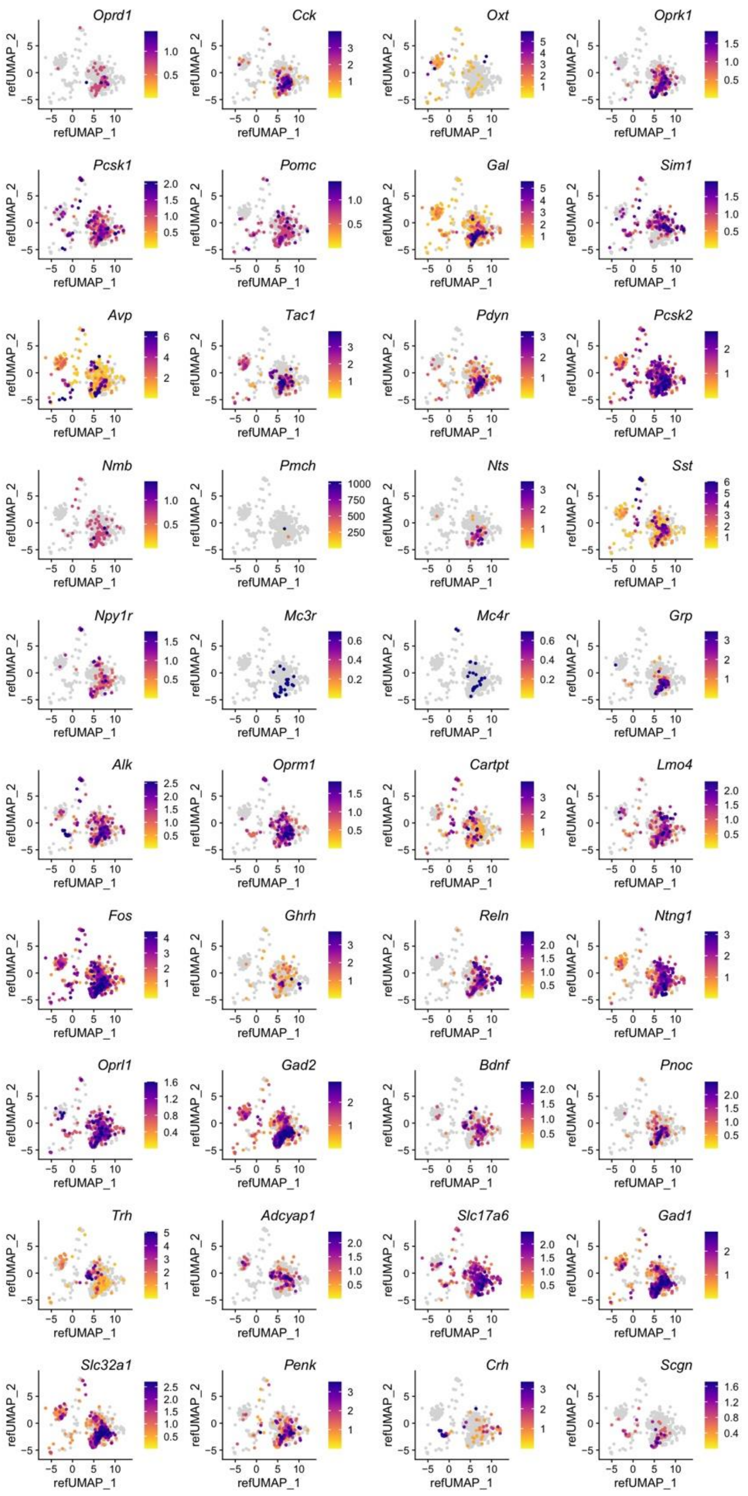


**Supporting Figure 3. Expression landscape of neuropeptides and with 10x data mapped onto an integrated Smart-seq2 reference.** UMAP plots show the expression of genes selected from 10x data[43] and mapped onto an integrated reference UMAP from Xu *et al.*[25].

Gueissaz *et al*. - Supporting Figure 4

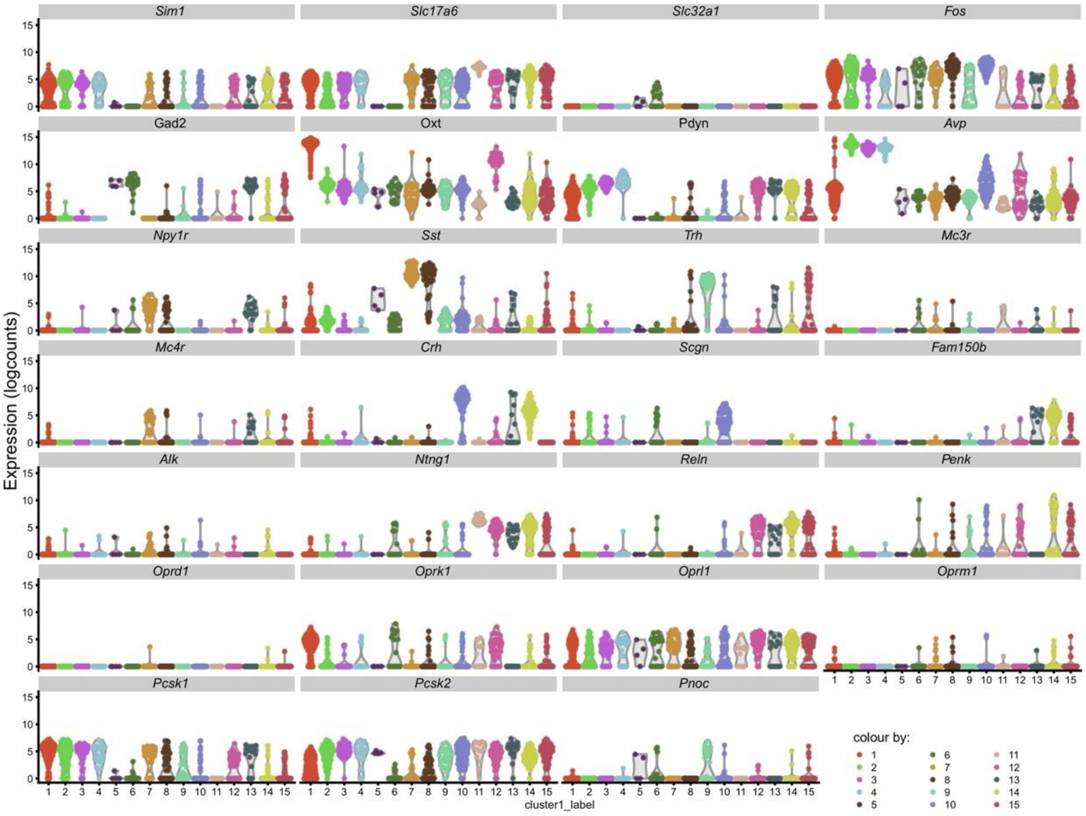


**Supporting Figure 4. Quantification of gene expression across neuronal clusters in the PVN.** Violin plots illustrate the distribution of normalized expression levels (logcounts) for selected genes across cell clusters (not sorted) identified in the PVN by Smart-seq2[25]. Each violin plot shows the density distribution of expression for a specific gene within a given cluster, providing a quantitative comparison of gene expression profiles across neuronal subpopulations.

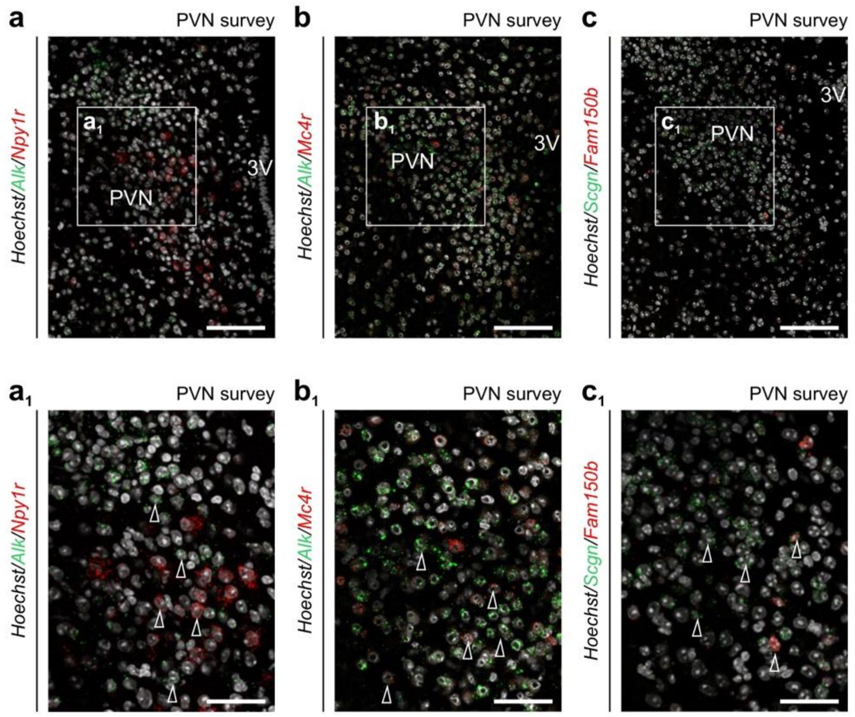


**Supporting Figure 5. Colocalization of *Alk* and *Fam150b* with other cellular markers in the PVN.** Multiple-labelling *in situ* hybridization for *Alk* and either *Npy1r* (**a**) or *Mc4r* (**b**) in the PVN. In both cases, *Alk* did not seem to co-localize with *Npy1r* or *Mc4r*, two receptors implicated in the control of metabolism and feeding behaviors[20–24]. (**c**) Multiple-labelling *in situ* hybridization for *Fam150b* and *Scgn* in the PVN. *Fam150b* and *Scgn* did not co-localize in the PVN, confirming data from single-cell RNA-seq[25]. *Scale bars* = 300 µm (*overviews*) and 30 µm (*insets*).
